# T606-phosphorylation deprives the function of Kaiso as a transcription and oncogenic factor

**DOI:** 10.1101/2020.03.23.003509

**Authors:** Wei Tian, Hongfan Yuan, Sisi Qin, Wensu Liu, Baozhen Zhang, Liankun Gu, Jing Zhou, Dajun Deng

## Abstract

It is well known that Kaiso protein encoded by *ZBTB33* gene is a transcription repressor and that Kaiso–P120ctn interaction increases the shift of Kaiso from the nucleus into the cytoplasm. However, the regulatory mechanisms of Kaiso compartmentalization are far from clear. Here, we reported that AKT1 could phosphorylate 606-threonine residue (<u>T</u>606) within the RSS<u>T</u>IP motif of Kaiso in the cytoplasm. The T606-phosphorylated Kaiso (pT606-Kaiso) could directly bind to 14-3-3 family proteins and the depletion of T606 phosphorylation by T606A mutation abolished most of the Kaiso–14-3-3 binding. In addition, the Kaiso–P120ctn interaction was essential for the pT606-Kaiso accumulation in the cytoplasm. Notably, enforced *14-3-3σ* (*SFN*) overexpression could increase the pT606-Kaiso accumulation in the cytoplasm and de-repress the transcription of Kaiso target gene *CDH1*. Decreased amounts of both pT606-Kaiso and CDH1 proteins were frequently observed in human gastric cancer tissues relative to paired normal controls. The mRNA levels of *14-3-3σ* and Kaiso target gene *CDH1* were positively and significantly correlated with each other in bioinformatics analyses using publicly available RNA-seq datasets for human normal tissues (*n*=11688, *r*=0.60, *p*<0.001) in the GTEx project and for cancer cell lines (*n*=1156, *r*=0.41, *p*<0.001) in the CCLE project. Furthermore, Kaiso T606A mutant (unable to be phosphorylated) significantly increased the migration and invasion of cancer cells *in vitro* as well as boosted the growth of these cells *in vivo*. In conclusion, Kaiso could be phosphorylated by AKT1 at the T606 and the pT606-Kaiso accumulates in the cytoplasm through binding to 14-3-3/P120ctn that de-represses the expression of Kaiso target gene *CDH1* in normal tissues. Decreased Kaiso phosphorylation may contribute to the development of gastrointestinal cancer. The status of Kaiso phosphorylation is a determinant factor for the role of Kaiso in the development of cancer.

## INTRODUCTION

Kaiso protein encoded by the *ZBTB33* gene (chrX:119,384,607-119,392,251) is a classic transcription repressor containing a zinc-finger domain and a BTB/POZ domain (1). The zinc-finger domain of Kaiso can bind to both methylated CGCG-containing and non-methylated Kaiso binding-specific sequences of Kaiso target genes, and the BTB/POZ domain can further recruit the complex of NCoR1 corepressor and histone deacetylases to target genes and repress their transcription in the nucleus (2–4). Recent studies show that Kaiso may also act as a transcription activator in the promoter context-dependent manner (5, 6).

As a transcription repressor, Kaiso controls the cell cycle through repressing *CCND1* and *CCNE1* expression, affects Notch signaling pathway in intestinal cells through targeting *DLL1* and *JAG1* promoter, and inhibits the proliferation and invasion of tumor cells through downregulating *MMP7*, *MTA2* and other genes (7–10). Kaiso also represses *CDH1* and *CDKN2A* expression (11, 12). Interestingly, it has been reported that the amount of nuclear Kaiso, but not total Kaiso, is correlated with the invasion or prognosis of cancers (13, 14). Kaiso-deficient mice show resistance to intestinal cancer (15). Apparently, the expression and subcellular location states of Kaiso determine its normal functions and roles in cancer development.

Kaiso is also a cytoplasm protein, which regulates WNT-related pathway through interacting with P120ctn (CTNND1) protein (Fig. S1) (1, 16). Differences of subcellular locations of Kaiso are observed between cultured cells and tissues (12, 17). The P120ctn-Kaiso complexes could shift from the nucleus to the cytoplasm (18). Kaiso nuclear-cytoplasmic trafficking could be affected by environmental factors, such as cigarette smoke, through MUC1 and P120ctn binding (19). However, detailed regulation machinery for the compartmentalization of Kaiso remains far from clear.

The 14-3-3 proteins are originally identified in the brain (20). There are seven human 14-3-3 isoforms (α/β, ε, η, δ/γ, τ, ζ, σ). These 14-3-3 isoforms are homologous proteins with approximately 50% amino acid identity, capable of forming either homo- or hetero-dimers (21–24). Recent findings have implicated 14-3-3 proteins as a key regulator of signal transduction events (25). Among the family, 14-3-3γ and 14-3-3σ (SFN) have been confirmed to play important roles in cancer development (26–29).

In the present study, we found, for the first time, that Kaiso could be phosphorylated at Thr-606 (<u>T</u>606) within the RSS<u>T</u>IP motif by the protein serine-threonine kinase AKT1 and T606-phosphorylated Kaiso (pT606-Kaiso) could efficiently interact with 14-3-3 family member and P120ctn proteins, and accumulate in the cytoplasm that in turn blocks transcription repressor function of Kaiso and de-repressed the expression of its target gene *CDH1*. Relative to the paired normal control tissues, the level of pT606-Kaiso was markedly decreased in human gastric tissues that could lead to transcriptional repression of the *CDH1* gene.

## MATERIALS and METHODS

### Cell lines and culture

The gastric cancer cell lines MGC803, BGC823, and SGC7901 were kindly provided by Dr. Yang Ke at Peking University Cancer Hospital; MKN45 cell line was purchased from the National Infrastructure of Cell Line Resource (Beijing, China). The human embryonic kidney HEK293T cell line was kindly provided by Professor Yasuhito Yuasa at Tokyo Medical and Dental University; RKO cell line, by Dr. Guoren Deng, University of California. HEK293T cells was cultured in DMEM medium containing 10% FBS, and all the others were cultured in RPMI1640 medium containing 10% FBS and 100 U/mL penicillin/streptomycin (Life Technologies, Carlsbad, CA, USA) at 37 °C in a humidified incubator with 5% CO_2_. These cell lines were tested and authenticated by Beijing JianLian Gene Technology Co., before use. Short tandem repeat (STR) patterns were analyzed using GoldeneyeTM20A STR Identifier PCR Amplification Kit.

### Gastric carcinoma tissues and ethical issue

Gastric cancer tissues and the paired normal surgical margin tissues were collected from 12 patients at Peking University Cancer Hospital from 2000 to 2001 and stored at −80 °C. The Institutional Review Board of the Peking University Cancer Hospital approved the study. All of the patients provided written informed consent.

### Plasmids and reagents

The full-length Kaiso-coding sequence of the *ZBTB33* gene was amplified from human cDNA of MGC803 cells with a primer set (forward 5’-attaaactcgaggcatggagagtagaaaactga-3’ and reverse 5′-cgcttcgaattcgtttagtaagactctggtattat-3’), then inserted into *Xho*I and *Eco*RI sites of pEGFP-C1 vector to generate pEGFP-C1-Kaiso expression vector. pEGFP-C1-Kaiso-T606 mutants were obtained by mutation PCRs using a primer set (forward 5’-gatagatcaagcgctattcctgcaatg - 3’ and reverse 5’-cattgcaggaatagcgcttgatctatc-3’) for 606Thr → Ala (T606A) mutation. Plasmid pCMV-3Tag-2C-Kaiso was generated by inserting the full-length Kaiso-coding sequence into *BamH*I and *Eco*RI sites of pCMV-3Tag-2C vector. Plasmid pEBG-GST-Kaiso was generated by inserting the full-length Kaiso coding sequence into *BamH*I and *Not*I sites of pEBG vector.

pcDNA3.1-HA-AKT1 vector was purchased from Addgene (#9008, MA, USA); pEZ-M56-14-3-3γ-mCherry, pEZ-M56-14-3-3σ-mCherry vectors, from FulenGen (EX-T4084-M56, EX-T4084-M98-5, EX-C0507-M98, Guangzhou, China); pENTER-Flag-14-3-3 isoforms (α/β, ε, η, δ/γ, τ, ζ, σ) were purchased from Vigene Bioscience (CH867785, CH897212, CH845486, CH898602, CH878525, CH824520, CH890307, Shandong, China); Insulin (P3376, Beyotime, Shanghai, China), IL-6 (Cat. 200-06, Proteintech, NJ, USA), EGF (PHG6045, Thermo Fisher Scientific, MA, USA), MK2206 (HY-10358, MedChemExpress, NJ, USA) were also used in the study. *P120ctn*-specific siRNAs (#1: sense 5’-gaaugugaugguuuaguuuu-3’ and antisense 5’-aacuaaaccaucacauucuu-3’; #2: sense 5’-uagcugaccuccugacuaauu-3’ and antisense 5’-uuagucaggaggucagcuauu-3’; #3: sense 5’-ggaccuuacugaaguuauuuu-3’ and antisense 5’-aauaacuucaguaagguccuu-3’) were synthesized by Genepharma (Shanghai, China). The quantitative RT-PCR primer sequences for detection of the level of *CDH1* mRNA were: forward 5’-gaacgcattgccacatacac-3’ and reverse 5’-gaattcgggcttgttgtcat-3’ (Tm=58 °C). The *Alu* RNA was used as the reference (forward 5’-gaggctgaggcaggagaatcg-3’ and reverse 5’-gtcgcccaggctggagtg-3’, Tm=60°C), as previously described (30).

### Cell transfection

X-tremeGENE siRNA Transfection Reagent or X-tremeGENE HP DNA Transfection Reagent (Cat. 04476093001, Cat. 06366236001, Roche, Mannheim, Germany) were used in cell transfection with siRNAs against *p120ctn* (final concentration, 100 nM) or Kaiso or its mutant expression vectors (2 μg/well in 6 wells plate) following manufacturer's instructions. The efficiency of gene overexpression or knockdown was determined 48 or 72 hrs post transfection by Western blotting. For stable transfection, G418 was added into the medium to select consistent GFP-Kaiso expressing MGC803 cells (final concentration, 750 μg/mL). Flow sorting assay was performed with a FACS Calibur flow cytometer (BD Biosciences, Franklin Lakes, US) after 48 hrs cell transfection, then sorted transfected-SGC7901 cells were grown in 6-well plates under the G418 selection.

### Wound healing assay

All cells were seeding in 6-well plates (5 wells/treatment). After reaching 95–100% confluence, the wound healing assays were performed (31). A pipette tip was used to gently scratch the cell monolayer. After two washes with PBS, the cells were cultured with serum-free RPMI-1640 medium. Images of wound healing were captured at different times.

### Transwell assays

Transwell assays (3 wells/treatment) were performed to determine the migration and invasion of cancer cells (31). For the migration assay, all cells (2×10^4^ cells per chamber) were separately resuspended in 180 μL of serum-free RPMI-1640 medium and seeded in the upper chambers (8-μm pores; Corning Inc., Corning, NY). For the invasion assay, the upper chamber pre-coated with Matrigel (BD Biosciences, Franklin Lakes, NJ, USA) was used. Then, cells (4×10^4^ cells per chamber) were separately seeding in the upper chambers. After 24 to 48 hrs’ incubation, these chambers were fixed with 4% paraformaldehyde for 30 min and stained with 0.1% crystal violet. Images of migrating and invading cells were captured using a microscope (Leica DMI4000B, Milton Keynes, Bucks, UK).

### Cell proliferation assay

Cell counting kit-8 (CCK-8 Kit; C0037, Beyotime, Shanghai, China) was used to detect cell proliferation (32). Briefly, all cells at the density of 2×10^3^ cells per well were seeding in 96-well plate. Cell proliferation was assessed at 0, 24, 48, and 72 hrs by adding 10 μL of CCK-8 solution to each well. After 2 hrs’ incubation with CCK-8, the absorbance at 450 nm was quantified by a microplate reader (Tecan Infinite M200 PRO, Switzerland). The average value for these wells was calculated for each treatment and statistically compared with Student t-test.

### Xenografts in NOD-SCID mice (33)

Cells re-suspended in 0.15 mL PBS (1×10^7^ cells/mL) were inoculated subcutaneously into the bilateral inguinal of 6 week-old female NOD-SCID mice (9 mice/group, 1×10^6^ cells per injection; purchased from Beijing Huafukang Biotech). Mice were sacrificed on the 28th inoculation day, and xenografts were separated, weighted, and photographed.

### Subcellular fractionation and de-phosphorylation treatment (34, 35)

To prepare cytoplasmic and nuclear extracts, small pieces of tissue or cultured cells at 80% confluence were homogenized in ice-cold buffer CERI of Nuclear and Cytoplasmic Extraction Reagent (7883, Thermo Fisher, MA, USA) with CERII and 1 х EDTA-free Protease Inhibitor Cocktail (REF04693159001, Roche, Mannheim, Germany) according to the Instruction. Samples were then vortexed, incubated on ice for 20 min, and centrifuged at 14,000 *g* for 15 min at 4 °C. The supernatants were recovered to obtain the cytosolic extracts. The pellets were washed sequentially with CERI and then incubated in buffer NER on ice and vortexed 15 sec every 1o min. After 4 times of vortex, centrifuging at 14,000 *g* for 15 min, the extract was collected as nuclear proteins. The purities of cytoplasmic and nuclear extracts were respectively verified by probing with anti-β-TUBULIN and anti-LAMIN B antibodies.

For the calf intestinal alkaline phosphatase (CIAP)-catalyzed de-phosphorylation, the cytoplasmic and nuclear extracts were aliquoted into two centrifuge microtubes. 1 μL of CIAP (p4978, Merck, Darmstadt, Germany) (>10 U/μL) of protein was added into one aliquot (30 μL; ratio of protein extracts to 10 × CIAP buffer, 27:3) and incubated for 30 min at 37 °C. BSA was used as negative control in equal amounts of protein extracts in 1 × CIAP buffer.

### Immunoprecipitation (IP) and Western blotting

Antibodies for Kaiso (sc-365428, Santa Cruz, USA), P120ctn (66208-1, Proteintech, IL, USA), pan-14-3-3 (sc-629, Santa Cruz), 14-3-3 family (α/β, ε, η, δ/γ, τ, ζ, σ) kits (#9769, CST, USA), 14-3-3σ (sc-100638, Santa Cruz), Ser/Thr/Tyr phosphor-protein (ab15556, Abcam, UK), phosphorylated AKT substrate [(R/K)X(R/K)XX(pT/pS)] (pAKT-sub; #9611, Cell Signaling Technology), CDH1 (#3195, CST), LAMIN B1 (66095-1, Proteintech), β-TUBLIN (66240-1, Proteintech), HA (M20003, Abmart, Shanghai, China), FLAG (66008-2, Proteintech), GFP (NB100-1614, Novus, CO, USA), GST (66001-1, Proteintech), GAPDH (660004-1, Proteintech) were used in the IP and Western blotting analyses.

After being pre-cleared with protein A/G-coupled Sepharose beads (Cat. 11134515001 and 11243233001, Roche, Mannheim, Germany) for 2 hrs, the nuclear or cytoplasmic lysate was immunoprecipitated with mouse anti-Kaiso antibody or anti-Phosphoserine/threonine/tyrosine antibody plus protein A/G Sepharose for 8 hrs at 4 °C. Mouse IgG was used as a negative control. The precipitates were washed six times with lysis buffer, and boiled in 1 × loading buffer. Protein samples were resolved by SDS–PAGE, and electroblotted onto nitrocellulose membranes, which were blocked in 5% skim milk in PBST and probed with antibodies according to the Instruction Manual.

### Phos-tag SDS-PAGE assay

This is a modified SDS-PAGE method based on the novel Phos-tag (36), which can bind to phosphorylated proteins and decrease their migration speed. Thus, this assay is often used to distinguish dephosphorylated proteins from phosphorylated proteins on the case of that a phosphorylation-specific antibody is not available. 50 μM Phos-tag (final concentration, Phos-tag Acrylamide AAL-107, WAKO, Japan) and 100 μM MnCl_2_ were mixed to prepare SDS-PAGE gel.

### GST-Pull down(37)

The day after pEBG-GST-Kaiso transfection, MGC803 cells (in 10-cm dishes, eight dishes/group) were further transfected with pENTER-Flag-14-3-3 members (α/β, ε, η, δ/γ, τ, ζ, σ) or negative control vector, respectively. 48 hrs post transfection, MGC803 cells were harvested and used to prepare lysate with cell lysis buffer with 1× Protease Inhibitor Cocktail (REF04693159001, Roche, Mannheim, Germany). The lysate was incubated with Glutathione Sepharose beads (20 μL for one group, 17-0756-01, GE healthcare, Sweden) at 4 °C overnight. Beads were washed six times with 500 μL cell lysis buffer. After the last centrifuging, the supernatant was removed as clean as possible, and the pellet was suspended in 20 μL 1 ×SDS sample buffer. Antibodies for GST-Kaiso and Flag-14-3-3 family proteins were used to detect the pulled down precipitant.

### Co-immunoprecipitation (Co-IP)

After pre-cleared with protein A/G-coupled Sepharose beads for 2 hrs, the soluble proteins from whole cell lysate were immunoprecipitated with anti-Kaiso (sc-365428, Santa Cruz) or anti-GFP (ab290, Abcam) or other antibodies plus protein A/G Sepharose overnight at 4 °C. Mouse IgG or rabbit IgG was used as a negative control. The precipitates were washed six times with lysis buffer, and boiled after 1 × SDS loading buffer was added. Protein samples were resolved by SDS–PAGE, and electroblotted onto nitrocellulose membranes, which were blocked with 5% skim milk in PBST and probed with the interacted protein antibodies. Loading cell number ratio for Input lane to IP lanes was 1:100.

### Preparation of pT606-Kaiso-specific antibody

Anti-pT606-Kaiso polyclonal antibodies were raised in rabbits challenged with the synthesized phosphorylated peptide LSD**RSS<u>pT</u>IP**AM, a sequence corresponding to amino acids 600-610 of wildtype Kaiso (Kaiso-wt), absorbed with non-phosphorylated peptide LSD**RSS<u>T</u>IP**AM, and enriched by the phosphorylated peptide LSD**RSS<u>pT</u>IP**AM. Peptide synthesis and immunization of the animals were done by YouKe (Shanghai, China). The non-phos-peptide was also used to prepare the control polyclonal antibodies against total Kaiso at the same time. The specificity of pT606-Kaiso and control antibodies against the corresponding peptide (LSDRSSpTIPAM or LSDRSSTIPAM) was detected with ELISA assay.

### ELISA analyses for polyclonal antibodies against pT606-Kaiso and total Kaiso

Polystyrene plates were coated with 1 μg/mL synthetic phosphorylated peptide LSDRSSpTIPAM or non-phosphorylated peptide LSDRSSTIPAM link-coupled by bovine serum albumin (BSA) in 1 × CBS buffer overnight at 4 °C, respectively, and were washed three times with PBS containing 0.05% Tween 20. Unbinding sites were blocked with 5% milk at room temperature for 2 hrs. The purified antibodies were added (100 μL/well) and incubated at 37 °C for 1 h r. After being washed with 0.05% Tween/PBS, plates were added HRP-labeled goat anti-Rabbit IgG (100 μL/well) and incubated at room temperature for 30 min. Peroxidase activity was measured with 0.15 mg/mL TMB substrate solution (100 μL/well). After 15 min at room temperature, the reaction was stopped by 2M H_2_SO_4_ (50 μL/well). Optical density absorbance (OD) at 450 nm was determined using a model 550 microplate reader.

### Immunohistochemical staining (IHC)

Rabbit pT606-Kaiso polyclonal antibody (1:100) and mouse monoclonal antibody for total Kaiso (1:50, sc-23871, Santa Cruz) were used in the IHC analysis. The REAL™ EnVision™ Detection system including anti-rabbit/mouse IgG secondary antibodies and peroxidase/diaminobenzidine (Dako, Agilent Technologies) was used to visualize the primary antibody‑binding cells according to the manufacturer’s protocol. Briefly, paraffin sections (4 μm) were dewaxed and rehydrated in xylene and ethanol. For antigen retrieval these sections were autoclaved for 3 min in 10 mM sodium citrate buffer containing 0.05% Tween-20 (pH 6.0) for pT606-Kaiso and in 1 mM EDTA buffer (pH 8.0) for total Kaiso. Then, these sections were immersed in 3% H_2_O_2_ for 10 min to block endogenous peroxidase. Following submerging in 5% bovine serum albumin (BSA) (cat. no. A1933; Sigma Life Science; Merck KGaA) for 60 min, the sections were incubated with the primary antibody overnight at 4 °C. The PBS washed sections were then treated with the REAL™ EnVision™ Detection system and counterstained with hematoxylin (0.125%; ZhongShan Jinqiao Biotechnology) at room temperature for 1 min. Normal rabbit IgG and normal mouse IgG (cat. no. ZDR 5003 and cat. no. ZDR 5006; ZhongShan Jinqiao Biotechnology) were used as negative controls. The former was diluted and incubated as for the pT606-Kaiso antibody, and the latter was diluted and incubated as for total Kaiso antibody).

### Confocal analysis

For mCherry-14-3-3, direct fluorescence was detected with laser confocal microscope assay. MGC803 cells with mCherry-14-3-3 overexpression were rinsed for three times with PBS, fixed with 1% paraformaldehyde in PBS 30 min at 37 °C, punched with 0.5% triton X-100 for 10 min at 37 °C, incubated with respective primary and secondary antibodies, washed for three times in PBS, counterstained with DAPI (1 μg/mL) for 5 min, and then examined with Leica SP5 Laser Scanning Confocal Microscopy. The antibodies for pT606-Kaiso (1 μg/μL, 1:100), for total Kaiso (commercial antibody (0.2 μg/μL, 1:20, sc-365428, Santa Cruz), and for P120ctn (1:100, 66208-1, Proteintech) were used as the primary antibodies; the FITC-labeled antibody against rabbit IgG (1:100, ab6717, Abcam) and CY5-labeled antibody against mouse IgG (1:100, Cat. No. 072-02-18-06, KPL Gaithersburg, MD, USA) were used as secondary antibodies for observation under Leica SP5 Laser Scanning Confocal Microscope and were analyzed with ImageXpress Micro high content screening System.

For the detection of Kaiso in tissues, fresh cryostat sections (4 μm) from gastric carcinoma and the paired normal tissues were fixed with 4% paraformaldehyde for 10 min at 37°C and treated with 0.5% triton X-100 for 10 min at 37 °C, and then incubated with respective primary, secondary antibodies, DAPI, and then examined as described above.

### Publicly available RNA-Seq, cDNA array, and other datasets

The RNA sequencing datasets in Cancer Cell Line Encyclopeida (CCLE) and Genotype-Tissue Expression (GTEx) projects were downloaded from official websites (portals.broadinstitute.org/ccle and www.gtexportal.org) (38–40). All raw data were transferred to Transcripts per kilobase of exon model per Million mapped reads (TPM) using uniform names for each protein-coding gene in the human genome. The Pearson correlation coefficient (r) of genes expression was calculated by R statistical software (version 3.6.1) with a set of procedures (41). Briefly, raw data was read through “read.table()” function, and an array was established to store data by “array()” function. Then the Pearson correlation coefficient was calculated by “cor()” function, and finally saving the data to files in csv format by the function of “write.table()”.

## RESULTS

### Discovery of phosphorylation of Kaiso in the cytoplasm

Transcription repressor Kaiso is also a cytoplasmic protein. The regulatory mechanism of Kaiso compartmentalization is still unknown. Previous study showed there might be putative phosphorylation sites within Kaiso in proteomic mass spectrometry analysis (Fig. S1) (42–44). We wondered if the putative phosphorylation affected Kaiso compartmentalization. In Phos-tag SDS-PAGE analyses, a Phos-tag could bind specifically to a phosphate group in proteins via metal ions, such as Zn^2+^ or Mn^2+^, which could be used to separate phosphorylated proteins from non-phosphorylated proteins (36). Thus, the Phos-tag assay was preliminarily utilized to analyze the phosphorylation status of Kaiso in human cells. We found that while the cytoplasmic and nuclear Kaiso from GFP-Kaiso stably transfected MGC803 cells migrated at the same speed in regular SDS-PAGE gel, the cytoplasmic Kaiso migrated much slower than the nuclear Kaiso in the Phos-tag gel, whether these cells were cultured *in vitro* or transplanted into nude mice as a xenograft, and no phosphorylated Kaiso was detected with the CIAP de-phosphorylation (Fig. 1A).

**Figure 1.**
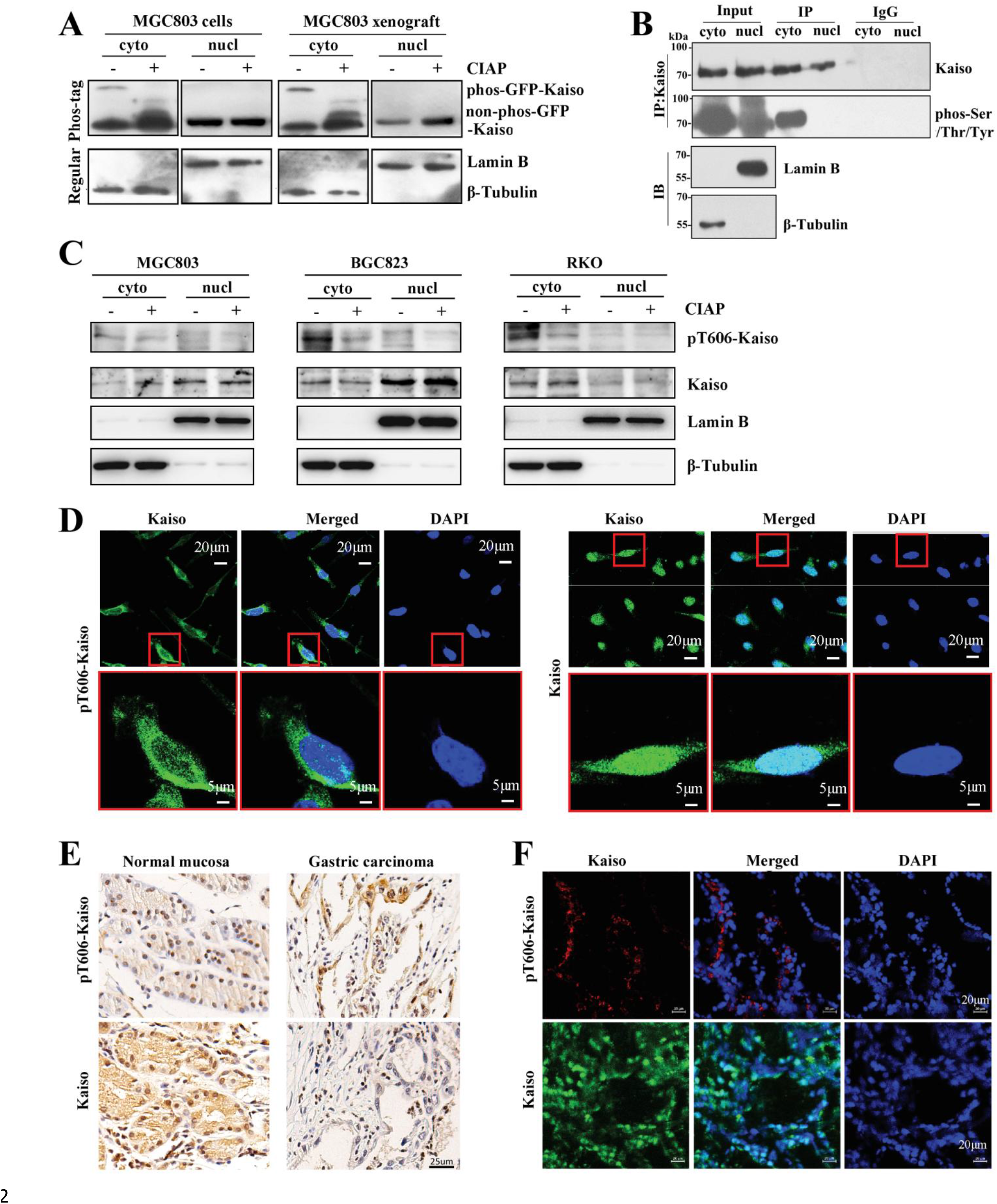
Different phosphorylation states of Kaiso protein in the cytoplasm and nucleus of cells *in vitro* and *in vivo*. (**A**) The phosphorylation statuses of GFP-Kaiso stably transfected MGC803 cells and the corresponding xenograft tissues in the Phos-tag SDS-PAGE analysis. (**B**) The phosphorylation statuses of endogenous cytoplasmic and nuclear Kaiso in MGC803 cells were validated using anti-phosphoSer/Thr/Tyr universal antibody. (**C**) Subcellular location of endogenous pT606-Kaiso and total Kaiso in cytoplasmic and nuclear proteins (with and without de-phosphorylation treatment by CIAP for 30 min) extracted from 3 cancer cell lines in Western blotting. (**D**) Locations of pT606-Kaiso and total Kaiso in the cytoplasm and nucleus in an indirect immunofluorescence confocal analysis. (**E**) Locations of pT606-Kaiso and total Kaiso in a representative gastric carcinoma and the paired normal tissues in IHC analysis. (**F**) Locations of pT606-Kaiso and total Kaiso in human gastric tissues in the indirect immunofluorescence confocal analysis.

To validate the different phosphorylation states of Kaiso in the cytoplasm and nucleus, an anti-phosphoserine/threonine/tyrosine universal antibody was then used to precipitate global phosphorylated proteins and a Kaiso-specific antibody was used to visualize the possible phosphorylated Kaiso (Fig. 1B). Again, phosphorylated Kaiso was observed only in the cytoplasmic precipitates, but not in the nuclear counterpart, suggesting that there may be phosphorylation of Kaiso in the cytoplasm. Thus, Kaiso phosphorylation kinase and target site were characterized in details as described below.

### Kaiso is phosphorylated at T606 by AKT1 kinase

Human Kaiso contains a conservative RSS<u>T</u>IP motif with the threonine-606 (T606) residue (Fig. 2A). Protein kinase B AKT1 is a typical kinase for the (RX)RXX<u>pS/pT</u> motif in multiple proteins such as mTOR, GSK-3β, AMPKA, Catenin-β1 (45–48). We wondered that the RSS<u>T</u>IP motif within Kaiso might be one of AKT1 targets. To verify if Kaiso T606 could be a true phosphorylation site for the kinase AKT1, we stimulated the activity of AKT signaling of MGC803 cells with insulin, IL-6, and FBS after overnight starvation (49). As expected, the phosphorylation level of endogenous Kaiso was markedly increased in MGC803 cells stimulated with insulin, IL-6, and FBS, but not with EGF (Fig. 2B).

**Figure 2.**
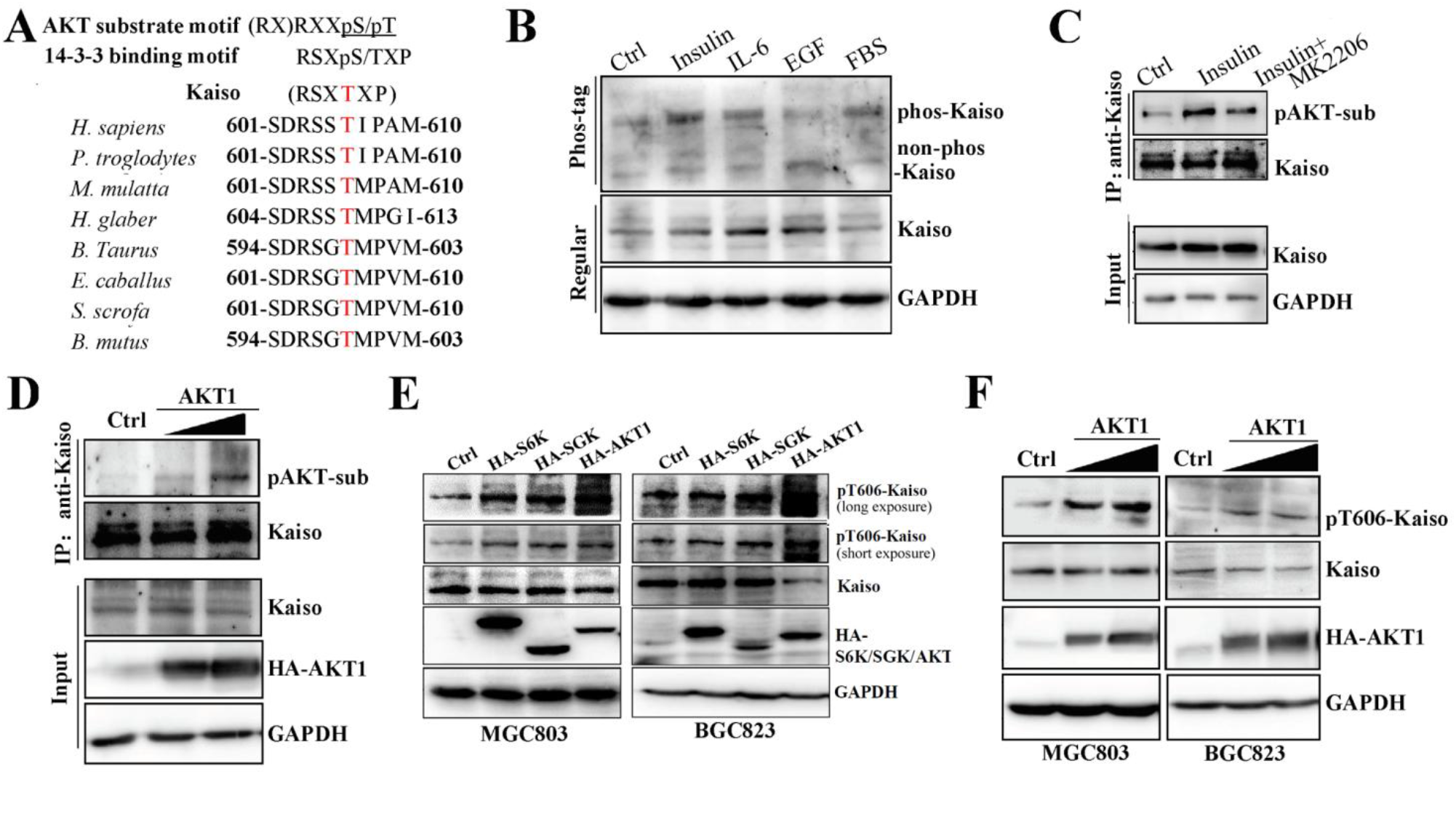
AKT1 increases the phosphorylation of Kaiso at T606. **(A**) A conservative RSX<u>T</u>XP motif within Kaiso of human and other species (**B**) After starvation overnight, treatments of Insulin (100 ng/mL), IL-6 (10 ng/mL) and fetal bovine serum (FBS, 1:10000 v/v) for 15 min increased the phosphorylation level of endogenous Kaiso in MGC803 cells. (**C**) Effects of AKT inhibitor MK2206 treatment (10 μmol/mL) for 30 min blocked the promotion of Insulin induced Kaiso phosphorylation as pAKT substrate in MGC803 cells. (**D**) *AKT1* overexpression at different doses increased the amount of phosphorylated AKT substrate (pAKT-sub) in Kaiso complexes immunoprecipitated by Kaiso antibody in MGC803 cells. (**E**) The activity comparison of three kinase candidates to phosphorylate Kaiso at T606 in MGC803 and BGC823 cells. (**F**) The T606-phosphorylation status of endogenous Kaiso in MGC803 and BGC823 with *AKT1* overexpression after starvation overnight.

Using the immunoprecipitation assay employing a phospho-(Ser/Thr) AKT substrate (pAKT-sub)-specific antibody, endogenous Kaiso signal could also be detected in the immunoprecipitated pAKT-sub protein complexes, and vice versa (Fig. S3A). The amounts of both global pAKT-sub and Kaiso-antibody-immunoprecipitated pAKT-sub were also significantly increased by insulin stimulation and reversed by AKT inhibitor MK2206 treatment (Fig. 2C). A similar effect was also observed in MGC803 cells with *AKT1* overexpression (Fig. 2D).

Then, a pT606-Kaiso-specific polyclonal antibody was prepared from rabbit using the phos-peptide LSD**RSS<u>pT</u>IP**AM as antigen (Fig. S2) and used to directly detect phosphorylated Kaiso. The activity of AKT1 to phosphorylate Kaiso at T606 was much higher than other two AGC kinases S6K and SGK (Fig. 2E). As expected, the amount of pT606-Kaiso was significantly increased in MGC803 and BGC823 cells with *AKT1* overexpression (Fig. 2F). Together, these results indicate that Kaiso can be phosphorylated at T606 by AKT1.

To confirm effect of the T606 phosphorylation on Kaiso compartmentalization, we further checked the phosphorylation status of endogenous Kaiso in the nucleus and cytoplasm using the pT606-Kaiso specific antibody. In consistent with the results described above, most pT606-Kaiso was detected in the cytoplasm of 3 human cancer cell lines (MGC803, BGC823, and RKO) and the CIAP de-phosphorylation treatment markedly decreased the amount of pT606-Kaiso in the cytoplasm in Western blotting (Fig. 1C). The confocal microscopy results further confirmed the cytoplasmic compartmentalization of pT606-Kaiso in MGC803 cells (Fig. 1D). Similarly, pT606-Kaiso was only detected in the cytoplasm in the IHC and confocal microscopy analyses while total Kaiso was mainly detected in the nucleus of human gastric mucosal tissues (Fig. 1E and 1F). Collectively, the above results demonstrate that AKT1 can phosphorylate Kaiso at T606 and pT606-Kaiso mainly locate in the cytoplasm of human cells.

### pT606-Kaiso interacted with 14-3-3 family members

It is well known that RSXpSXP is a 14-3-3 phosphoserine binding consensus motif (50). To study whether the Kaiso RSS<u>pT</u>IP is a 14-3-3 binding motif, we performed GST-pull down and Co-IP experiments. The results of GST-pull down assay showed that GST-Kaiso could strongly pull down 14-3-3σ (SFN), moderately pull down 14-3-3ε, 14-3-3**γ**, 14-3-3ζ, and weakly pull down 14-3-3η in MGC803 cells lysate (Fig. 3A). The Co-IP results confirmed that endogenous Kaiso could bind to endogenous pan-14-3-3 or 14-3-3σ protein in MGC803 cells (Fig. 3B). As expected, the T606A mutation abolished most Kaiso-14-3-3 interaction (Fig. 3C), suggesting that the phosphorylation site T606 within the RSS<u>T</u>IP motif may play a main role in determining the Kaiso–14-3-3 interaction. In addition, AKT1 overexpression increased the Kaiso–14-3-3 interaction (Fig. S3B). These data indicate that Kaiso could interact with 14-3-3 family proteins in a T606 phosphorylation-dependent manner.

**Figure 3.**
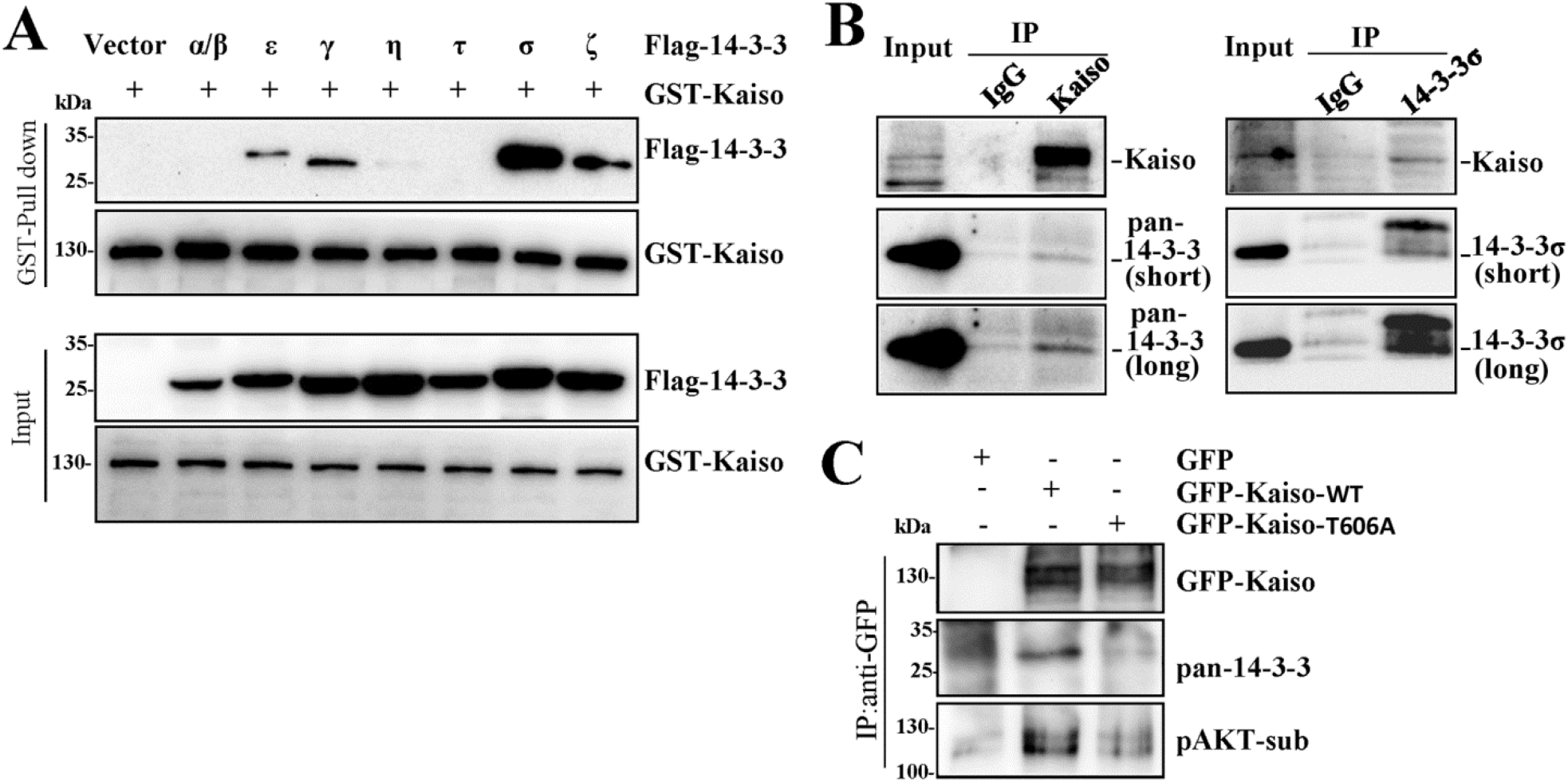
Kaiso interacted with 14-3-3 family members depending on Kaiso pT606. (**A**) After the plasmids of 14-3-3 family members and GST-Kaiso were co-transfected into MGC803 cells, GST-Kaiso pulled down various isoforms of 14-3-3 family, especially 14-3-3σ. (**B**) Endogenous Kaiso immunoprecipitated pan-14-3-3 or 14-3-3σ protein immunoprecipitated endogenous Kaiso in MGC803 cell in Co-IP analysis. (**C**) T606A mutation of Kaiso concealed its interaction with pan-14-3-3 in MGC803 cells in Co-IP analysis.

### Increased cytoplasmic accumulation of pT606-Kaiso by the 14-3-3σ interaction

Intracellular Kaiso compartmentalization is affected by growth conditions (13, 17). Our indirect immunofluorescence confocal microscope analyses showed that 81% endogenous Kaiso was observed in the nucleus of MGC803 cells (Fig.4A-B). Notably, enforced mCherry-14-3-3γ or −14-3-3σ overexpression significantly increased the proportion of endogenous Kaiso in the cytoplasm, compared to the mCherry control vector (from 19% to 37% or 36%) and Kaiso was mainly co-localized with mCherry-14-3-3γ or -14-3-3σ in the cytoplasm of these cells in the confocal analysis. Western blotting confirmed that mCherry-14-3-3σ overexpression increased the cytoplasmic accumulation of endogenous Kaiso (Fig. 4C, red arrowed) while it decreased the nucleic Kaiso (Fig. 4C, green arrowed). In addition, 14-3-3σ only promoted the cytoplasmic accumulation of wildtype Kaiso (Kaiso-wt), but not the T606A mutant (Fig. 4D, red arrowed), implying an increase of pT606-Kaiso in the cytoplasm by 14-3-3σ and likes. 14-3-3σ or 14-3-3γ overexpression also induced more cytoplasmic pT606-Kaiso in MGC803 and BGC823 cell lines in Western blotting (Fig. S3C-D). These results solidly confirm that the pT606-Kaiso can interact with 14-3-3 and in turn accumulate in the cytoplasm of human cancer cells.

**Figure 4.**
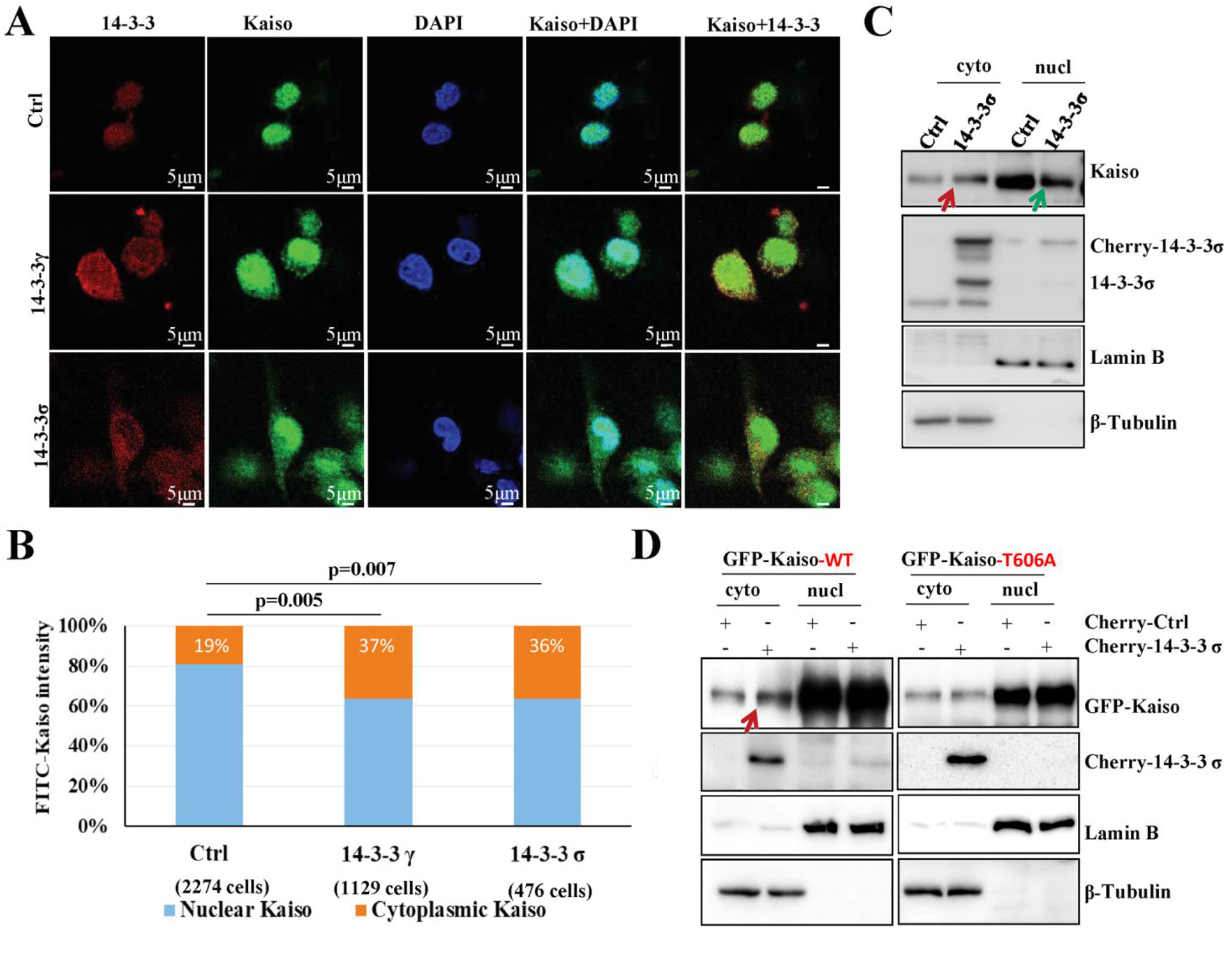
14-3-3 proteins promote the cytoplasmic accumulation of Kaiso in Kaiso T606 phosphorylation-dependent manner. (**A**) The subcellular location of endogenous Kaiso in MGC803 cells with or without 14-3-3γ or 14-3-3σ overexpression by indirect immunofluorescence staining assay. (**B**) Proportion of Kaiso in the nucleus and in cytoplasm of MGC803 cells with and without 14-3-3γ or 14-3-3σ overexpression. (**C**) Western blot for detecting the amounts of endogenous Kaiso in cytoplasm and nucleus protein in MGC803 cells with 14-3-3σ overexpression. (**D**) Comparison of the levels of GFP-Kaiso (wild type) or T606A mutant in the cytoplasm and nucleus in MGC803 cell with and without 14-3-3σ overexpression.

### The P120ctn interaction is essential for pT606-Kaiso-14-3-3 accumulation in the cytoplasm

As a given Kaiso interacting protein, the P120ctn binding is essential for the trafficking of Kaiso from the nucleus to the cytoplasm (18, 19). Thus, we further studied whether 14-3-3 bound to the pT606-Kaiso–P120ctn complex or individual pT606-Kaiso molecule in the cytoplasm. Interestingly, we found that 14-3-3σ overexpression markedly increased the amount of P120ctn in the anti-Kaiso antibody precipitated complexes, while the levels of total endogenous Kaiso and P120ctn proteins were not changed in MGC803 cells (Fig. 5A). Further analysis showed that more Kaiso-P120ctn binding was detected in the cytoplasm, but not in the nucleus of MGC803 cells with 14-3-3σ overexpression (Fig. 5B). Immunofluorescence confocal microscopy showed that P120ctn, Kaiso, and 14-3-3σ were mainly co-localized in the cytoplasm (Fig. 5C). Moreover, when P120ctn was knocked down by siRNA (siP120ctn), the cytoplasmic Kaiso accumulation promoted by 14-3-3σ disappeared in the MGC 803 cells (Fig. 5D) while the level of total Kaiso was not changed (Fig. 5E). These results suggest that the P120ctn interaction is essential for the pT606-Kaiso–14-3-3σ accumulation in the cytoplasm.

**Figure 5.**
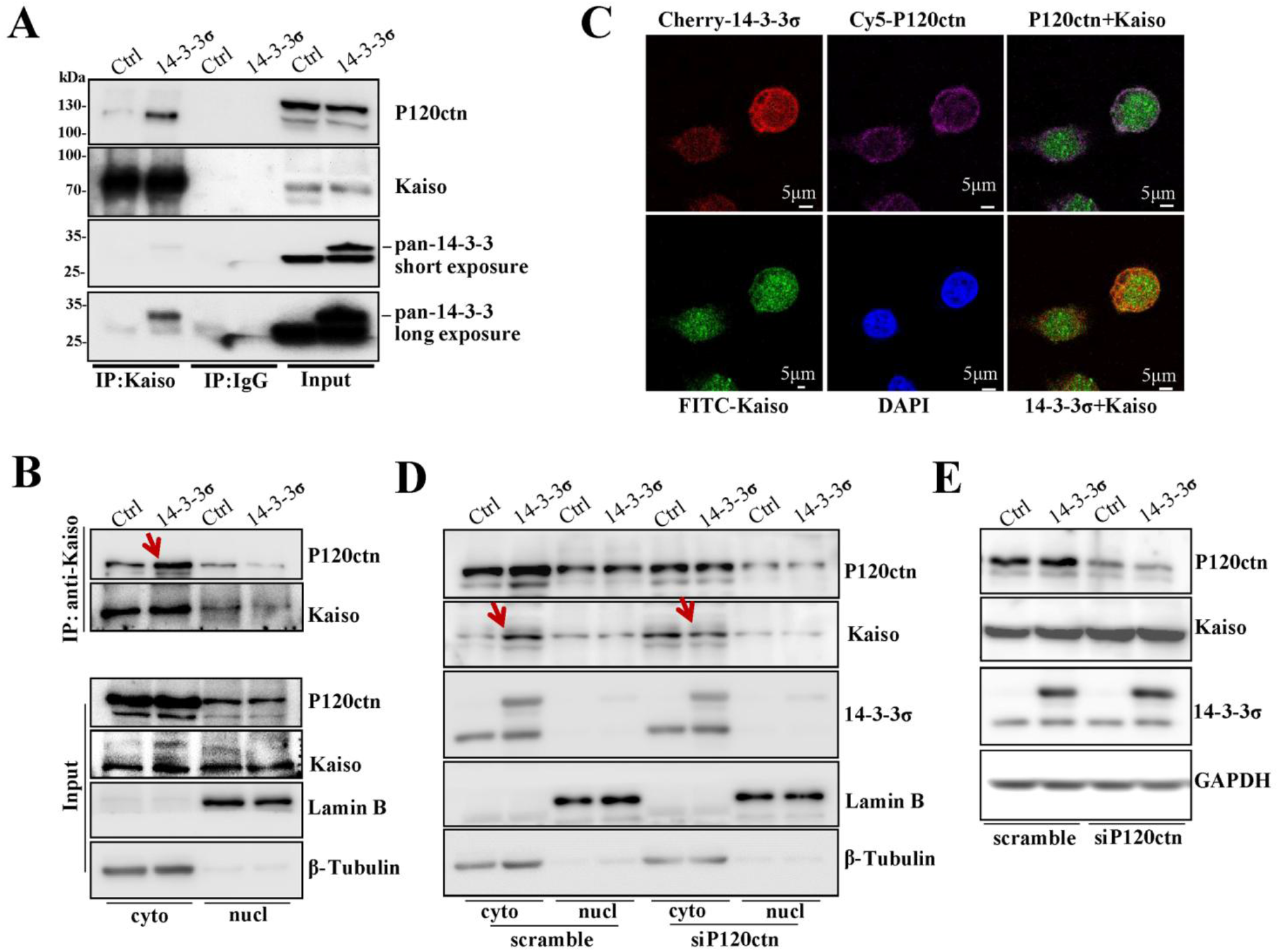
14-3-3σ promotes the interaction of Kaiso and p120ctn in the cytoplasm. (**A**) More Kaiso-p120 complex was immunoprecipitated by Kaiso antibody in MGC803 cells with 14-3-3σ overexpression in Co-IP assay. Total protein levels of p120ctn and 14-3-3σ in the lysate of MGC803 cells with 14-3-3σ overexpression were displayed in the right side. (**B**) Alterations of interactions between Kaiso and p120ctn proteins in the cytoplasm and nucleus of these cells with *14-3-3σ* overexpression in the Co-IP assay using Kaiso antibody. (**C**) The subcellular locations of Kaiso, p120ctn, and mCherry-14-3-3σ in MGC803 cells. (**D**) Western blotting images for detecting effect of *p120ctn* knockdown and *14-3-3σ* overexpression on distribution of Kaiso in the cytoplasm and nucleus of MGC803 cells. (**E**) The levels of total p120ctn and Kaiso proteins in the lysate of MGC803 cells with *14-3-3σ* overexpression and siRNAs knockdown of *p120ctn* expression.

### De-repression of Kaiso target genes by T606 phosphorylation

It was reported that Kaiso could bind to both methylated and non-methylated Kaiso binding sequences in target gene promoters and suppress their transcription (11). We wondered whether T606 phosphorylation of Kaiso and its bindings with 14-3-3/P120ctn proteins could affect expression of Kaiso target genes through the cytoplasmic accumulation. Through re-analyzing the RNA-seq datasets for 14-3-3 family members and Kaiso target genes (including *CDH1*, *CCND1*, *CCNE1, MTA2, DLL1,* and *DAG1*) from GTEx and CCLE (38-40), we found that the level of 14-3-3σ mRNA was positively and most significantly correlated with that of *CDH1* in both normal human tissues (*n*=11688, *r*=0.60, *P*<0.001 in Pearson correlation analysis) in GTEx datasets and cancer cell lines in CCLE datasets (*n*=1156, *r*=0.41, *P*<0.001) (Fig. 6A and Fig. S4A). This is consistent with the phenomenon that 14-3-3σ could strongly bind to Kaiso described above (Fig.3A), suggesting an exact effect of 14-3-3σ expression on regulation of *CDH1* transcription, probably through the pT606-Kaiso–14-3-3σ interaction. Interestingly, association between the levels of *CDH1* and *Kaiso* mRNA was only observed in human normal tissue samples from subjects (*n*=570) in the GTEx and most normal tissue samples patients (*n*=697) in the Cancer Genome Atlas (TCGA) datasets, but not in cancer cell lines (*n*=1063) in the CCLE datasets nor in cancer tissues of many organs in the TCGA datasets (Fig. S4B and S4C), suggesting that endogenous Kaiso may be not a repressor for *CDH1* transcription in normal cells via the translocation of phosphorylated Kaiso from the nucleus to the cytoplasm, and may be a repressor for *CDH1* transcription in cancer cells because of blocks of the nucleus-cytoplasm shift by the decreased level of Kaiso phosphorylation as described below.

**Figure 6.**
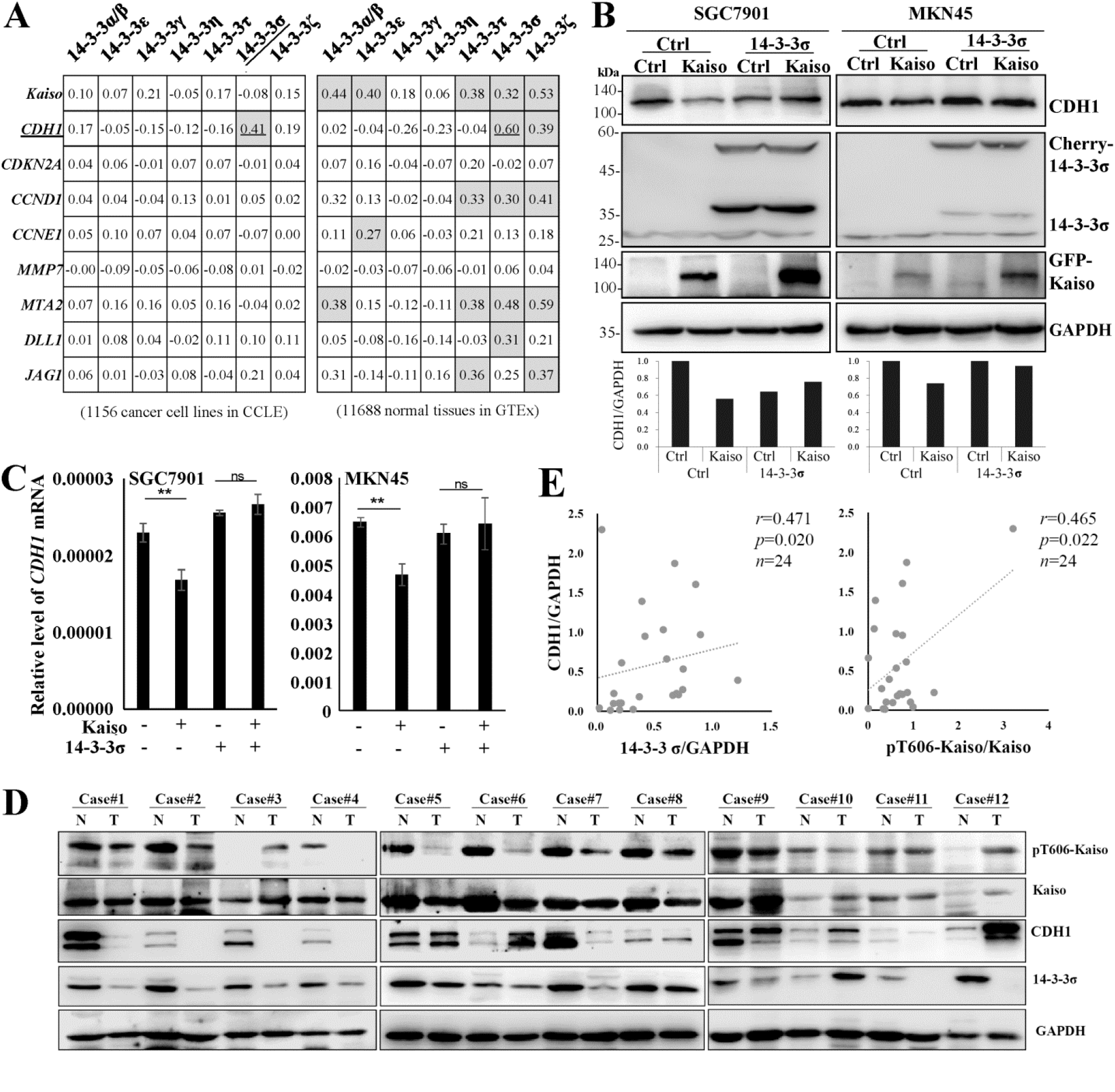
Effects of 14-3-3σ on inhibition of Kaiso’s target gene *CDH1* expression. (**A**) The Pearson correlation coefficient between mRNA levels of 14-3-3 family members and Kaisos’ target genes in public RNA-seq datasets from Cancer Cell Line Encyclopedia (CCLE, 1156 cancer cell lines) and Genotype-Tissue Expression (GTEx, 11688 human normal tissues). (**B**) The level of *CDH1* in SGC7901 and MKN45 cells with or without wild type Kaiso and 14-3-3σ overexpression. (**C**) The level of the *CDH1* mRNA in SGC7901 and MKN45 cells with or without wild type Kaiso and 14-3-3σ overexpression (**D**) The amounts of pT606-Kaiso, total Kaiso, 14-3-3 σ, and CDH1 proteins in gastric carcinoma (T) and the paired normal tissues (N) from 12 patients by Western blotting. (**E**) Correlation between ratios of pT606-Kaiso to total Kaiso and CDH1 to GAPDH proteins based on the density of these proteins in Western blot.

**Figure 7.**
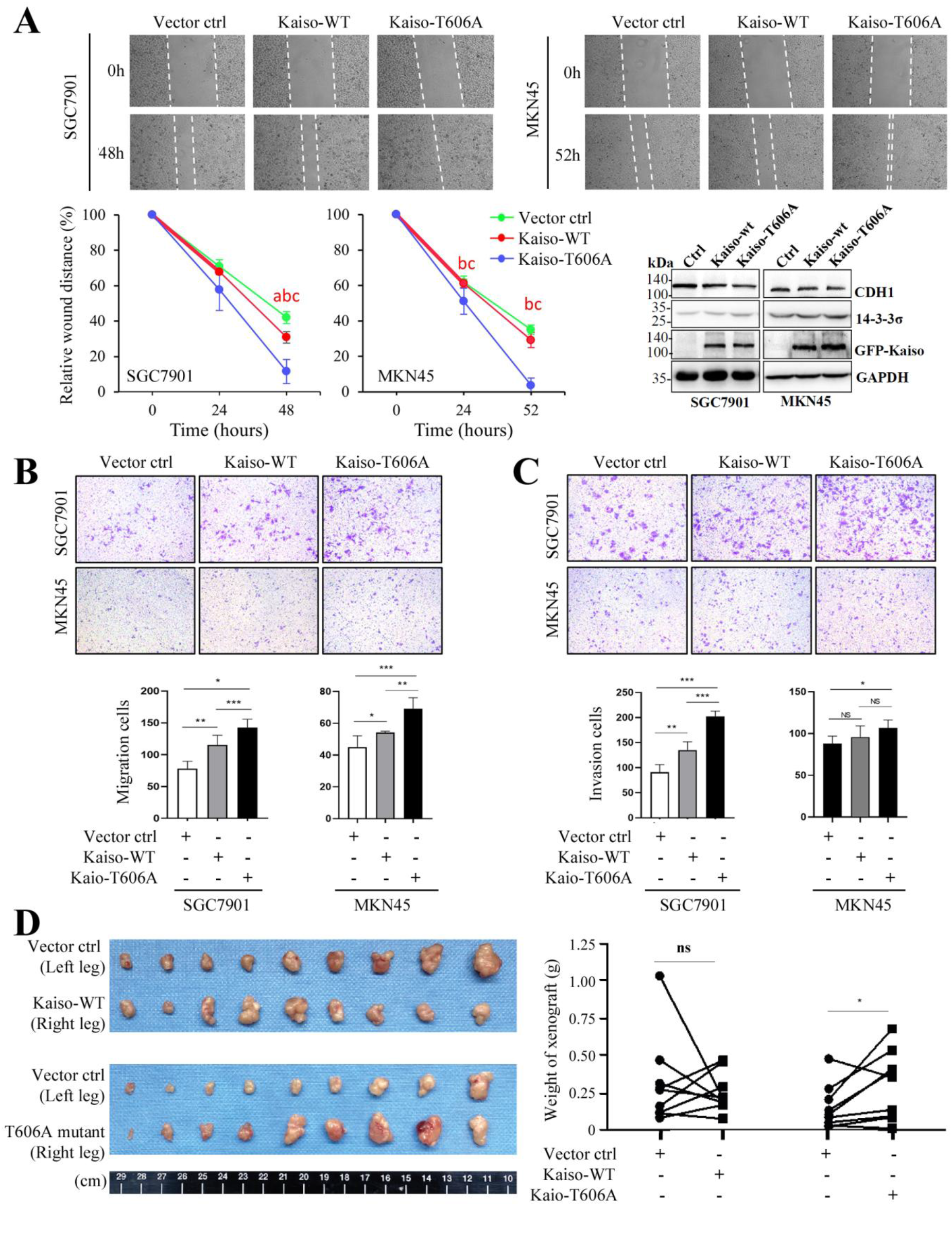
Effect of wildtype Kaiso and its T606A mutant on gastric cancer cell migration and invasion in vitro and growth in vivo. (**A**) The percentage of open wounds was calculated to determine the migration of gastric cancer (GC) SGC7901 and MKN45 cells overexpressing Kaiso-WT or its T606A mutant (a/b/c: Kaiso-WT vs. Ctrl/Kaiso-T606A vs. Ctrl/Kaiso-T606A vs. Kaiso-wt, p < 0.05); The results of Western blot were also inserted to illustrate the status of Kaiso overexpression; (**B** and **C**) The results of transwell assays to show the migration and invasion of GC cells overexpressing Kaiso-WT or Kaiso-T606A mutant, respectively; (**D**) Tumors derived from SGC7901 cells transfected with Kaiso-WT and its T606A mutant or empty vector control. */**/***: p<0.05/0.01/0.001

Because we did not detected *CDH1* expression in MGC803 and BGC823 cells, therefore, two gastric cancer cell lines SGC7901 and MKN45 with active *CDH1* expression were used to study effects of Kaiso and 14-3-3σ expression changes on the function of Kaiso as a transcription repressor. As expectedly, Kaiso overexpression alone indeed decreased the amount of CDH1 whereas 14-3-3σ co-overexpression could abolish Kaiso-induced *CDH1* repression in SGC7901 and MKN45 cells in Western blotting (Fig. 6B). These results were confirmed in qRT-PCR analyses (Fig. 6C). A similar relationship was observed in gastric carcinoma tissues. While the level of pT606-Kaiso was decreased in most gastric cancer samples, the ratios of pT606-Kaiso to total Kaiso was positively and significantly associated with the amounts of CDH1 (adjusted by GAPDH) among gastric cancer and the paired normal tissue samples from 12 patients (Fig. 6D and 6E). Similar correlation was also detected between the levels of CDH1 and 14-3-3σ. Taken together, the above results indicate that pT606-Kaiso-14-3-3σ binding in the cytoplasm can deprive the function of Kaiso as a transcription repressor and lead to de-repression of the transcription of Kaiso target gene *CDH1*.

### Enhancement of the growth of cancer cells by depriving pT606 phosphorylation of Kaiso

To evaluate the impact of pT606 phosphorylation of Kaiso on biological behaviors of cancer cells, we performed a set of functional assays through overexpression of wildtype Kaiso or its T606A mutant that cannot be phosphorylated. The results of both wound healing and transwell tests showed that the migration of SGC7901 and MKN45 gastric cancer cells was mostly increased by transient transfection of Kaiso T606A mutant while it was only weakly increased by the transfection of wildtype Kaiso (Fig. 8A and 8B). Similar differences were also observed in the invasion test (Fig. 8C). Although difference was not observed in the proliferation of cells (data not shown), however, the animal experiments showed that the average weight of xenografts derived from SGC7901 cells stably transfected with Kaiso-T606A mutant was significantly higher than that of cells transfected with empty vector control in NOD-SCID mice (Fig. 8D, *p*<0.05 in the paired t-test). All these results showed that Kaiso T606A mutant significantly increased the migration and invasion of cancer cells *in vitro* as well as boosted the growth of cancer cells *in vivo* compared to Kaiso T606 wildtype.

## Discussions

The subcellular locations of Kaiso determine its roles in normal cell differentiation and cancer development. However, detailed regulation machinery for the compartmentalization of Kaiso is far from clear. In this study we demonstrated, for the first time, that Kaiso could be phosphorylated at T606 by AKT1, the pT606-Kaiso could interact with 14-3-3 and P120ctn in the cytoplasm and promote the shift of Kaiso from the nucleus to the cytoplasm. The phosphorylation of Kaiso finally leads to deprivation of Kaiso as a transcription repressor and de-repression of Kaiso target gene *CDH1* in normal tissues. Depletion of the T606 phosphorylation of Kaiso could augment repression of *CDH1* expression and promote the growth of cancer cells *in vivo*.

It’s well known that 14-3-3 family proteins bind to common phosphoserine/phosphothreonine-containing peptide motifs corresponding to Mode-1 (RSXpSXP) or Mode-2 (RXY/FXpSXP) sequences (46). We found that Kaiso contains a very conservative RSS<u>T</u>IP motif that could be phosphorylated by AKT1 in both *in vivo* and cell-free system at RSS<u>T</u>IP-T606 site. This is consistent with the report that T606 is one of phosphorylation sites of Kaiso in mass spectrometry analysis (42). That pT606-Kaiso could directly bind 14-3-3 family proteins, and the T606A mutation could abolish most Kaiso–14-3-3 binding indicate the T606 phosphorylation is essential for Kaiso–14-3-3 binding.

It has been reported that a region consisting of N-terminal amino acid residues (454 - 672aa) of Kaiso directly interacts with P120ctn (1) and Kaiso–P120ctn interaction promotes cytoplasmic-nuclear trafficking of Kaiso (18, 19). The phosphorylation site T606 is located within the P120ctn binding site, suggesting an effect of the phosphorylation at T606 on Kaiso–P120ctn interaction. In addition, P120ctn could bind to CDH1 and modulate its function and stability (51) and WNT-stimulated P120ctn phosphorylation could promote P120ctn releasing from the CDH1–P120ctn complexes, enhances Kaiso–P120ctn interaction (52). It is well known that most 14-3-3σ proteins localize in the cytoplasm (53) while Kaiso mainly localizes in the nucleus (54). Here, we observed that pT606-Kaiso, 14-3-3, and P120ctn proteins could co-localize in the cytoplasm and siRNA-knockdown of *P120ctn* abolished the 14-3-3-induced cytoplasmic Kaiso–14-3-3 binding. These results demonstrate that Kaiso–14-3-3σ interaction may be dependent on the Kaiso–P120ctn binding and that Kaiso, 14-3-3σ, and P120ctn may form a triplex in the cytoplasm. It was reported that cytoplasm Kaiso functionally linked the autophagy-related protein LC3A/B in breast cancer cells (55). How Kaiso, 14-3-3σ, and P120ctn interacting with each other is worth further studying.

Human *Kaiso/ZBTB33* gene locates in chromosome X, which is frequently amplified in the genome of many cancers. It is controversy on Kaiso’s roles in cancer development. In the absence of the tumor suppressor *APC*, *Kaiso*-deficient mice were resistant to intestinal cancer, suggesting that *Kaiso* might be an oncogene (15). On the contrary, Kaiso has also been suggested to be a potential tumor suppressor, which repressed the transcription of *MMP7*, *CCND1*, and *WNT11* genes involved in oncogenesis and metastasis (7, 9, 56). Functions of Kaiso are tightly related and significantly influenced by microenvironmental factors (17). We found here that increase of the level of pT606-Kaiso or pan-14-3-3 (especially 14-3-3σ) expression could lead to Kaiso accumulation in the cytoplasm, deprive transcriptional function of Kaiso, and de-repression of Kaiso target gene *CDH1*.

Kaiso phosphorylation may account for the positive correlation between the levels of *ZBTB33/Kaiso* and *CDH1* mRNAs in human normal tissues. In contrast, the decreased level or deprivation of pT606 phosphorylation of Kaiso could block the cytoplasmic transportation of Kaiso, repress *CDH1* transcription, and promote the growth of cancer cells. Our results are consistent with the report that overexpression of *Kaiso/Zbtb33* resulted in downregulation of *Cdh1* in mice intestinal tissues (57). These phenomena indicate that Kaiso phosphorylation may be a crucial determinant for functions of Kaiso in cancer development through de-repression of tumor related genes.

In conclusion, Kaiso can be phosphorylated by AKT1 at T606 within the very conservative motif RSS<u>T</u>IP in the cytoplasm. pT606-Kaiso can directly interact with 14-3-3 and P120ctn proteins and accumulate in the cytoplasm that consequently leads loss of functions of Kaiso as a transcription and cancer repressor. The decreased level of phosphorylation of Kaiso may contribute to cancer development through downregulation of *CDH1* expression in the stomach.

## Abbreviations

BSA: bovine serum albumin
CIAP: calf intestinal alkaline phosphatase
CY5: Cyanine5
DAPI: 4’, 6-diamidino-2-phenylindole
ELISA: enzyme-linked immunosorbent assay
FITC: fluorescein isothiocyanate
HC: Heavy chain of IgG
IP: immunoprecipitation
KBS: Kaiso binding sequence
NOD-SCID mice: Non-obese diabetic/severe combined immunodeficient mice
P120ctn: P120 catenin (CTNND1)
pAKT-Sub: phosphorylated AKT substrate
PBS: Phosphate-buffered saline
Phos-tag: Phos-tag molecular binds specifically to phosphate group in proteins via metal ions
pT606: T606-phosphorylated
T606A: replacement of the 606th amino acid residue threonine with alanine
wt: wildtype

## Acknowledgements

We gratefully acknowledge the critical reading of the manuscript and kindly sending AGC kinase plasmids by Dr. Jian-Ping Guo in Sun Yat-Sen University. This work is supported by National Natural Science Foundation of China (81402343).

## Competing Interest

The authors declare that they have no conflict of interest.

## Author contribution

D. Deng, W. Tian and S. Qin. designed research; W. Tian, H. Yuan, S. Qin, and B. Zhang performed research; D. Deng, W. Tian, H. Yuan, L. Gu and J. Zhou analyzed data; D. Deng, W. Tian and S. Qin wrote the paper.

---

**Figure S1.**
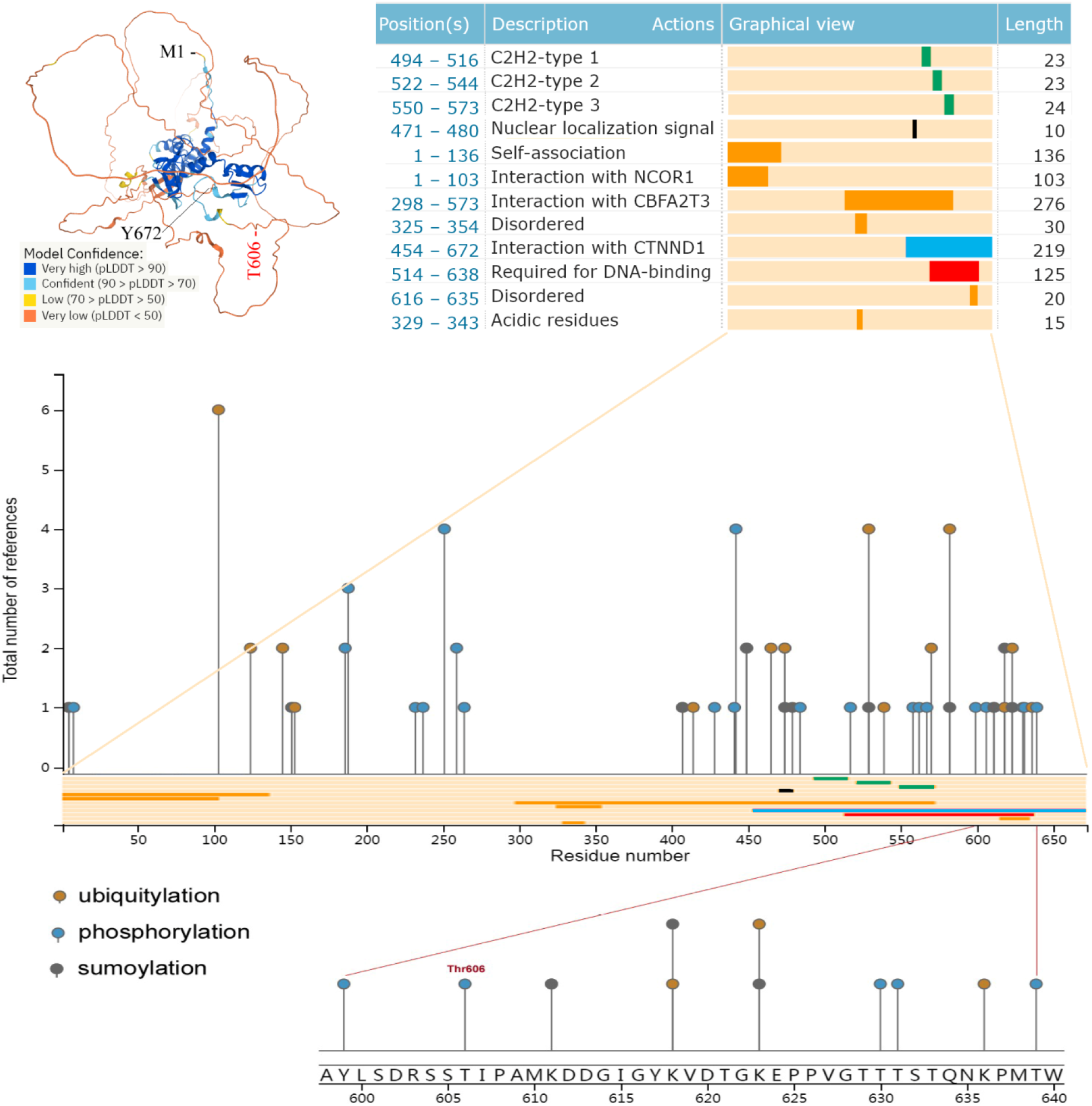
The status of Kaiso structure and protein modifications detected by LC/MS. Phosphorylation at the Thr606 residue in the LSDRSS<u>T</u>IPAM motif was illustrated in the bottom chart in details. The image was adapted with graphs for Kaiso modifications from the web site (www.phosphosite.org) (42, 43). AlphaFold predicted 3D structures for Kaiso (ZBTB33/Q86T24) protein was adapted from images downloaded from the website (https://alphafold.ebi.ac.uk) [44]; pLDDT, AlphaFold produced per-residue confidence score between 0 and 100. Information for Kaiso domains was adapted from images downloaded from the website (https://www.uniprot.org/uniprot/Q86T24) (1, 45).

**Figure S2.**
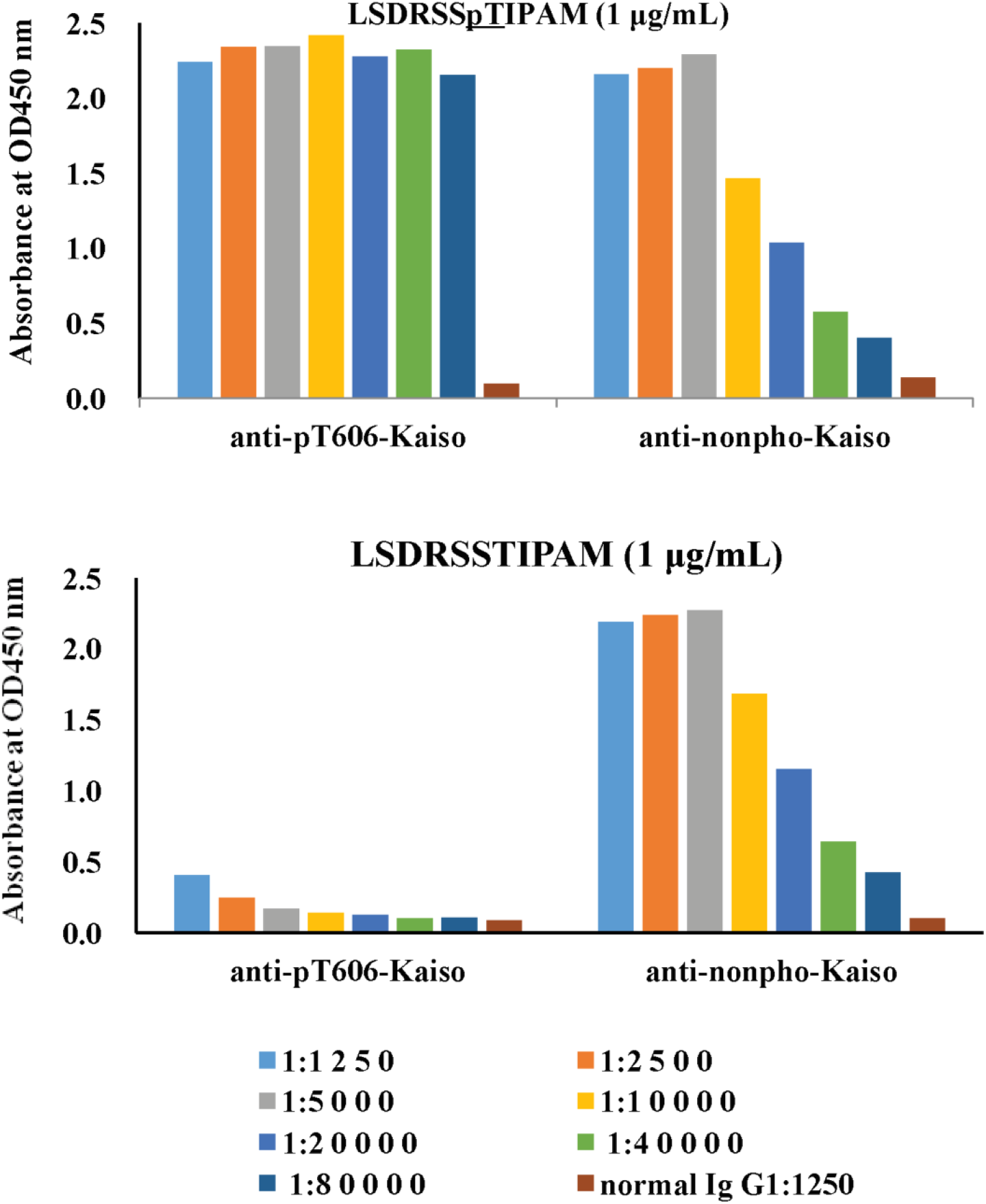
Characterizing the specificity of pT606-Kaiso polyclonal antibody. ELISA results for the specificity of pT606-Kaiso and control antibodies against the pT606-Kaiso peptide (LSDRSS<u>pT</u>IPAM, the top chart) and for the specificity of pT606-Kaiso and control antibodies for the nonphosphorylated control peptide (LSDRSS<u>T</u>IPAM, the bottom chart).

**Figure S3.**
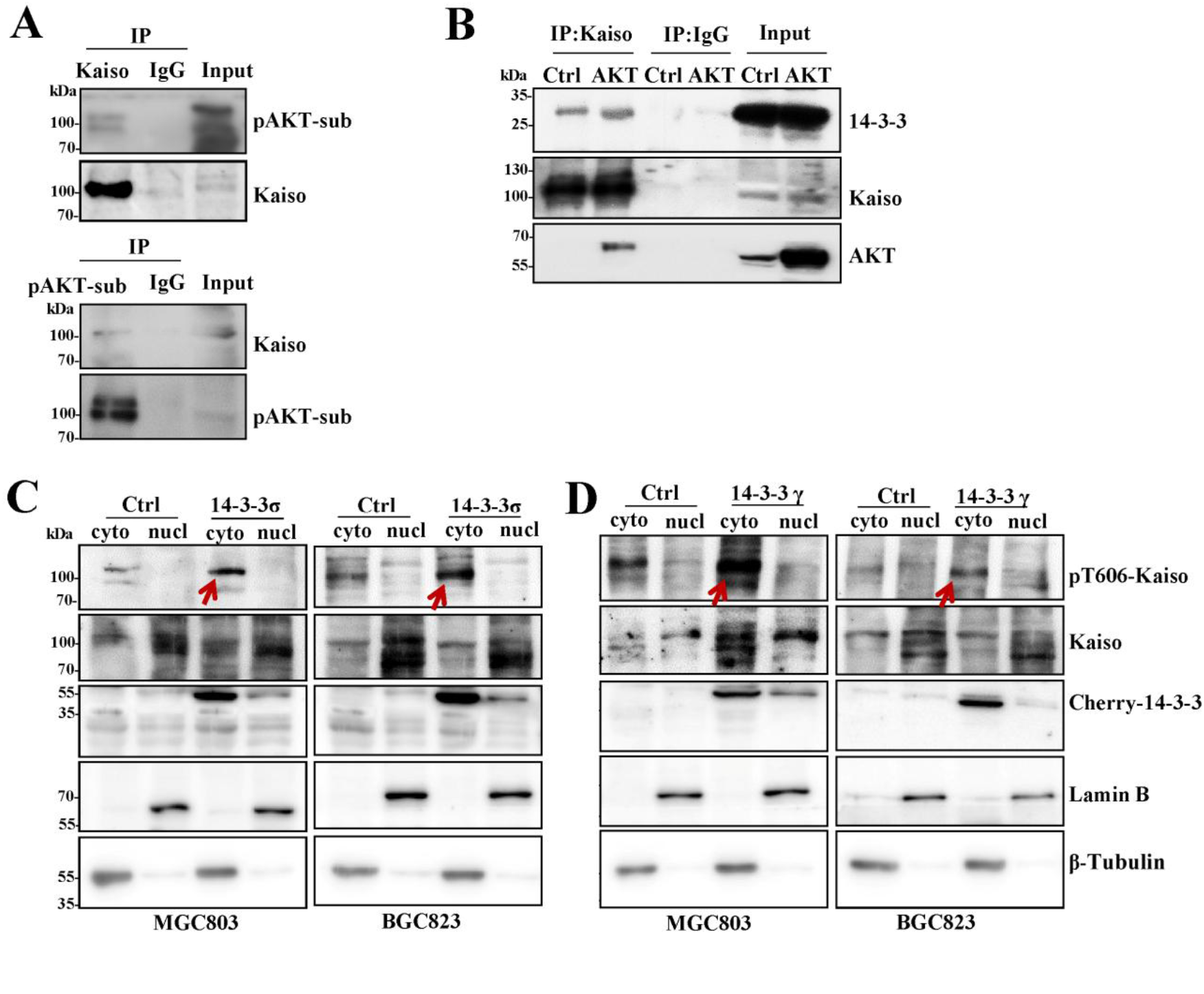
AKT1 and 14-3-3 regulate the T606-phosphorylation and subcellular localization of endogenous Kaiso. **(A**) Endogenous Kaiso in MGC803 cells immunoprecipitated by Kaiso antibody was identified by the antibody specific for AKT substrate motif, and the immunoprecipitation by AKT substrate antibody was identified by antibody against Kaiso. (**B**) *AKT1* overexpression increased endogenous Kaiso-14-3-3 interaction in MGC803 cells in Co-IP assay. (**C** and **D**) The T606-phosphorylation states of endogenous Kaiso in the cytoplasm and nucleus of MGC803 and BGC823 cells with 14-3-3σ and 14-3-3γ overexpression, respectively.

**Figure S4.**
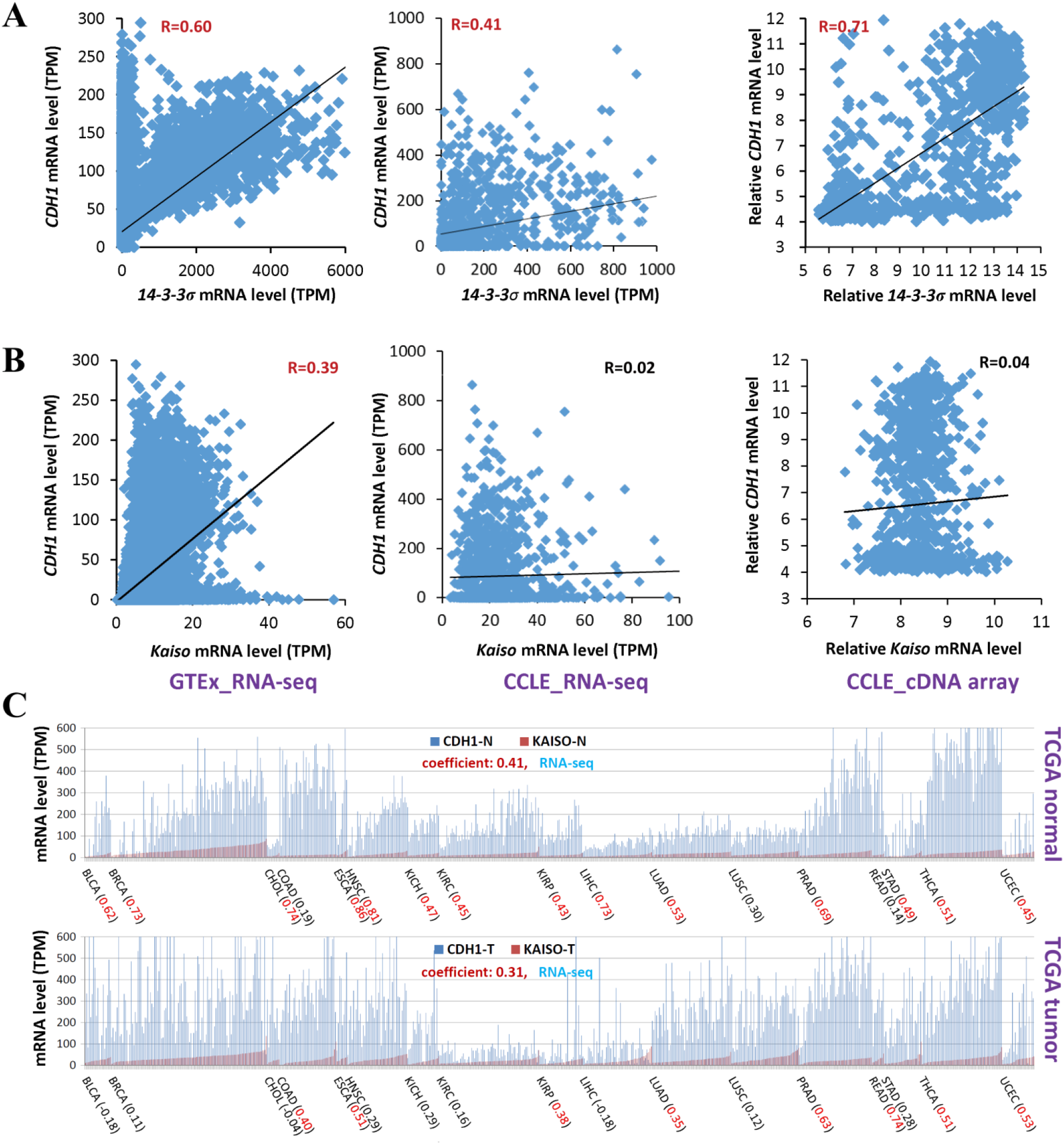
Correlation between the levels of *CDH1* and *14-3-3σ* or *Kaiso/ZBTB33* mRNAs in RNA-seq and cDNA array datasets. (**A** and **B**) Human normal tissues in the Genotype-Tissue Expression (GTEx) project and cancer cell lines in Cancer Cell Line Encyclopedia (CCLE) project; (**C**) Human tumor tissue and the paired normal tissue samples from patients in the Cancer Genome Atlas (TCGA) project. Gene expression coefficient is labeled within parentheses for each kind of tissues.

## Notes

### Competing Interest Statement

The authors have declared no competing interest.

### Summary of Updates

To evaluate the impact of phosphorylation of Kaiso on CDH1 expression and growth of cancer cells, we carried out additional experiments. For example, we confirmed the de-repression effect of pT606-Kaiso on CDH1 expression using gastric cancer and the paired normal tissue samples from 12 patients (revised Fig.6D and 6E). We also added the results of additional experiments on effects of pT606-Kaiso on the growth of cancer cells in vitro and in vivo (revised Fig.7). An extra author, Dr. Hongfan Yuan, was added into the author list because she did these supplemental experiments.

